## Supplementary Material for "Human embryoid bodies as a novel system for genomic studies of functionally diverse cell types"

Supplementary Materials and Methods

**Reference integration and label transfer**

To assess the quality of our reference integration strategy (see Materials and Methods), we asked whether 1) datasets are being over-corrected and 2) EB cells annotated using reference cell types express expected marker genes.

We first subsetted EB cells to broad cell type categories identified using clustering (at resolution 0.1) and differential expression analysis: Pluripotent (cluster 0), Early Ectoderm (cluster 1), Endoderm (cluster 4) , Mesoderm (clusters 2, 6) , Neural Crest (cluster 3), and Neurons (cluster 5). Using each subset of cells, we repeated the reference integration pipeline by merging the EB cells with three reference data sets (fetal cells, hESCs, and an external set of Day 20 EBs), normalizing using *SCTransform*, running PCA using 5000 variable features, integrated using Harmony, and transferred labels based on the 5 nearest reference cells (see Materials and Methods) (Butler et al. 2018; Hafemeister and Satija 2019; Korsunsky et al. 2019). We found that 79% of EB cells are assigned to the same cell type in the full integration and subset integration. Of EB cells that are annotated differently in the full and subset integrations, 82% were labelled as 'hESC' or 'uncertain' in either the full or subset integration. This suggests that differences in these annotations are often be due to slight changes in the positioning of cells between the hESC reference cells and fetal reference cells; this is expected when pluripotent cells are not included in subsets of EB cells. And, importantly, cells do not often get annotated as a different fetal cell type. Together, these results suggest that this integration approach is robust to subsetting input cell types and in likely not over-integrating the test and reference data sets.

Next we asked whether annotated EB cells differentially express expected marker genes. We limited this analysis to annotations with at least 10 EB cells total from at least 2 individuals in two replicates. We then calculated pseudobulk expression for cells of the same annotation, individual, and replicate, and filtered genes to include only those with at 10 counts in at least some samples and at least 15 counts total across all samples. We then TMM-normalized pseudobulk expression values, used *voom* to calculate a weighted gene expression value, and tested for differential expression between annotations using limma. Of the annotations tested, the most significantly differentially expressed genes often included known cell type markers. For example, cells annotated as cardiomyocytes showed significant upregulation of of MYL7, MYL4, and TNNT2 (Fig. S18). Cells annotated as hepatoblasts showed significant upregulation of AFP, FGB, and ACSS2 (Fig. S18). Cells annotated as mesothelial cells showed significant upregulation of NID2, collagen genes (COL6A3, COL1A1, COL3A1, COL6A1). These results provide further support that our reference integration approach yields meaningful annotation of EB cells (Fig. S18).

Supplementary Figures and Tables

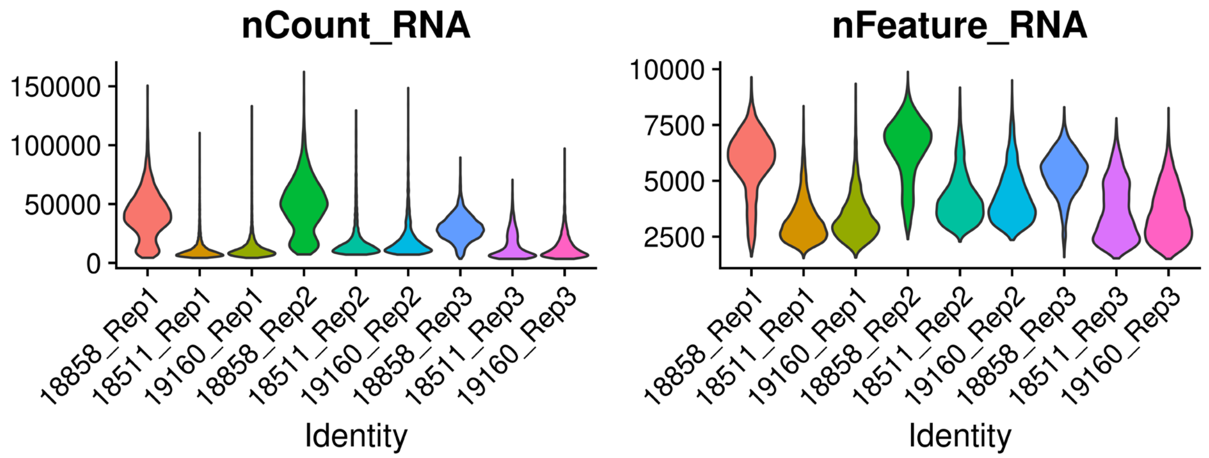


**Figure S1: Quality metrics after filtering.** Left) Violin Plot showing the total UMI counts in cells from each individual in each replicate after filtering. Right) Violin Plot showing the number of genes (features) expressed in cells of each individual and each replicate after filter.

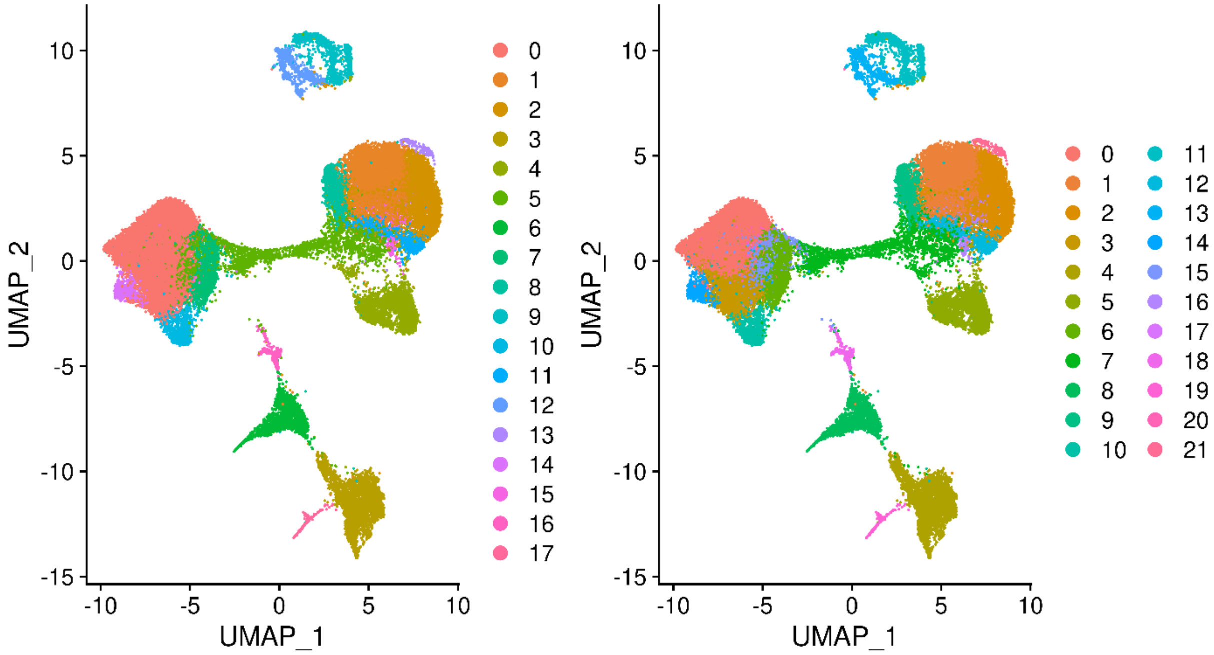


**Figure S2: Seurat Clusters identified at clustering resolution 0.5 (Left) and 0.8 (Right).**

56

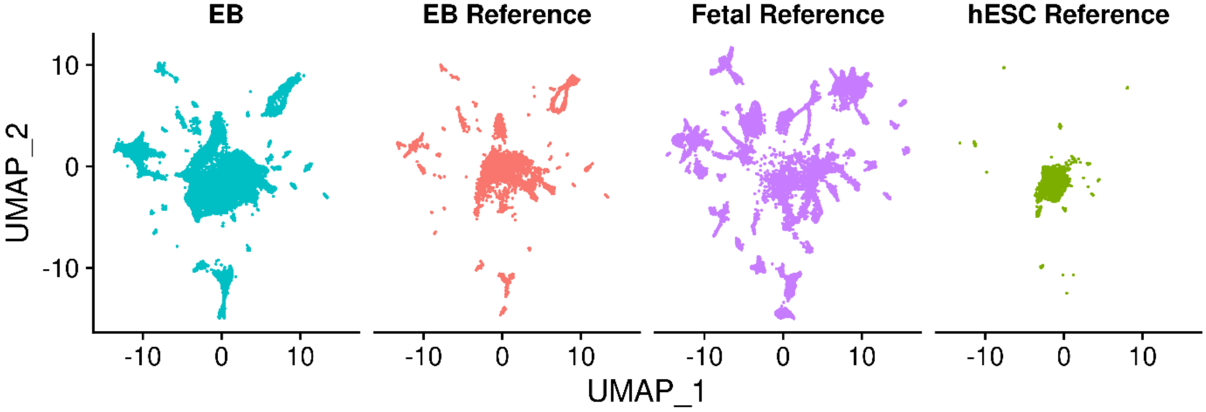


**Figure S3: UMAP visualization of EB cells and cell from each reference set after integration separated data set.**

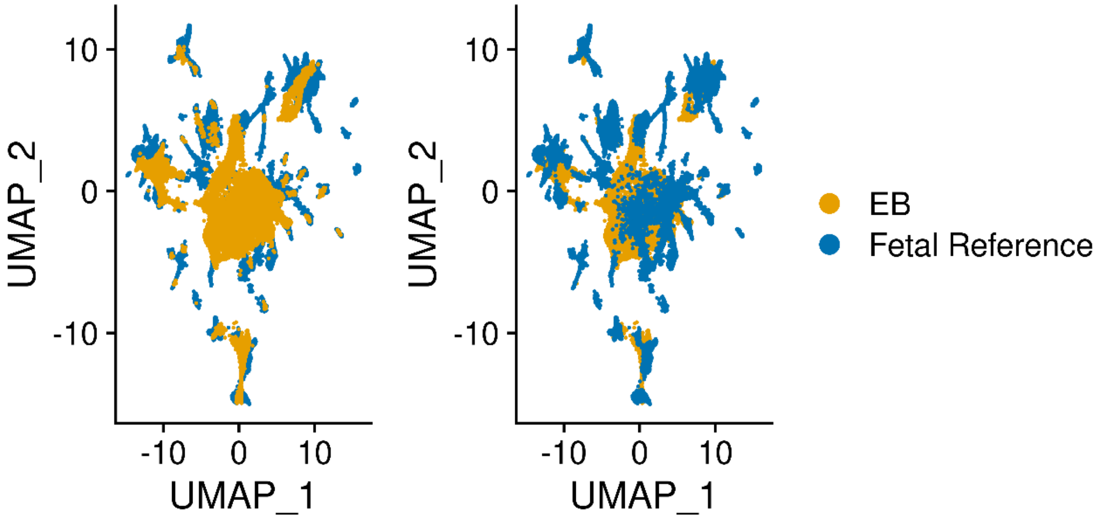


**Figure S4: UMAP visualization of EB cells from this study and fetal reference cells after integration. Cells are colored by data set.**

| Cell Type Annotation | Frequency in EB cells | Cell Type Annotation | Frequency in EB cells |
|---|---|---|---|
| Acinar cells | 52 | Lymphatic endothelial cells | 2 |
| Adrenocortical cells | 7 | Megakaryocytes | 37 |
| AFP_ALB positive cells | 36 | Mesangial cells | 222 |
| Amacrine cells | 79 | Mesothelial cells | 160 |
| Antigen presenting cells | 1 | Metanephric cells | 142 |
| Astrocytes | 39 | Microglia | 44 |
| Bipolar cells | 40 | MUC13_DMBT1 positive cells | 181 |
| Cardiomyocytes | 521 | Myeloid cells | 1 |
| CCL19_CCL21 positive cells | 2 | Neuroendocrine cells | 4 |
| Chromaffin cells | 24 | Oligodendrocytes | 53 |
| Ciliated epithelial cells | 314 | Parietal and chief cells | 14 |
| CLC_IL5RA positive cells | 3 | PDE11A_FAM19A2 positive cells | 35 |
| Corneal and conjunctival epithelial cells | 21 | Photoreceptor cells | 9 |
| Ductal cells | 329 | Purkinje neurons | 35 |
| ELF3_AGBL2 positive cells | 23 | Retinal pigment cells | 391 |
| Endocardial cells | 5 | Retinal progenitors and Muller glia | 21 |
| ENS glia | 30 | scHCL.hESC | 34086 |
| ENS neurons | 72 | Schwann cells | 38 |
| Epicardial fat cells | 112 | Skeletal muscle cells | 5 |
| Erythroblasts | 34 | SKOR2_NPSR1 positive cells | 39 |
| Excitatory neurons | 1 | SLC24A4_PEX5L positive cells | 297 |
| Ganglion cells | 61 | Smooth muscle cells | 60 |
| Goblet cells | 271 | Squamous epithelial cells | 160 |
| Granule neurons | 158 | Stellate cells | 247 |
| Hematopoietic stem cells | 9 | Stromal cells | 217 |
| Hepatoblasts | 212 | Syncytiotrophoblasts and villous cytotrophoblasts | 17 |
| Horizontal cells | 53 | Thymic epithelial cells | 116 |
| IGFBP1_DKK1 positive cells | 341 | Thymocytes | 1 |
| Inhibitory interneurons | 9 | Trophoblast giant cells | 5 |
| Inhibitory neurons | 65 | uncertain | 1566 |
| Intestinal epithelial cells | 501 | Unipolar brush cells | 37 |
| Islet endocrine cells | 552 | Ureteric bud cells | 26 |
| Lens fibre cells | 51 | Vascular endothelial cells | 58 |
| Limbic system neurons | 15 | Visceral neurons | 119 |

**Table S1: Frequency of each cell type present in EB data after transferring annotations from the fetal and hESC reference sets.**

59

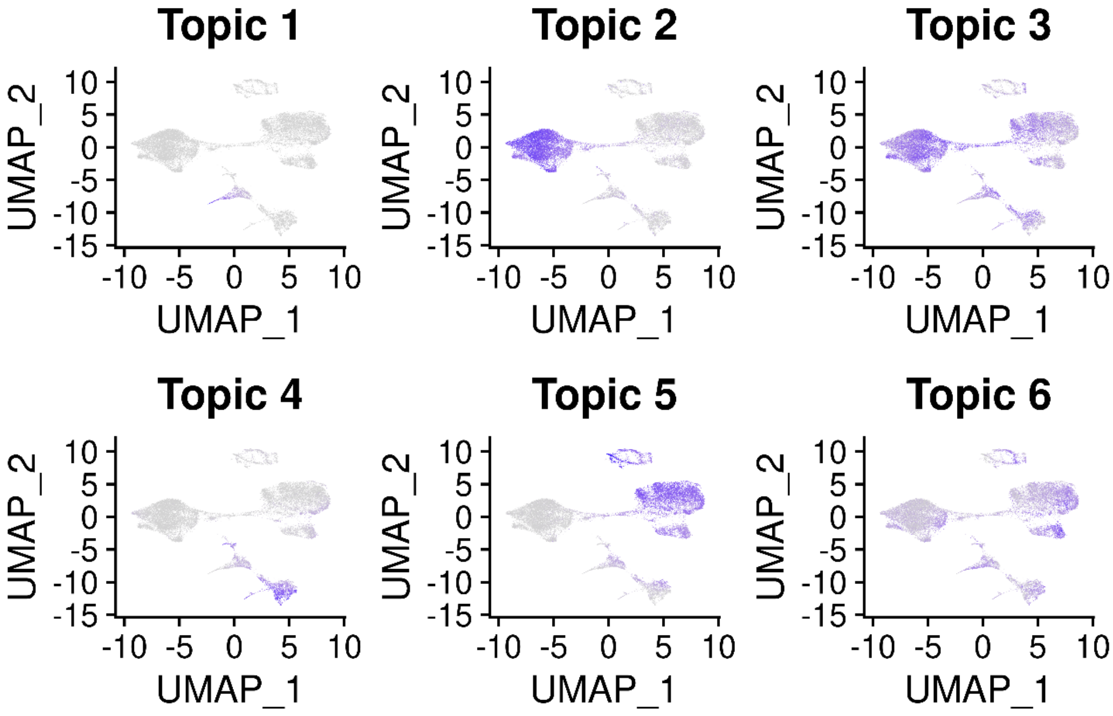


**Figure S5: UMAP visualization of k=6 topic loadings.**

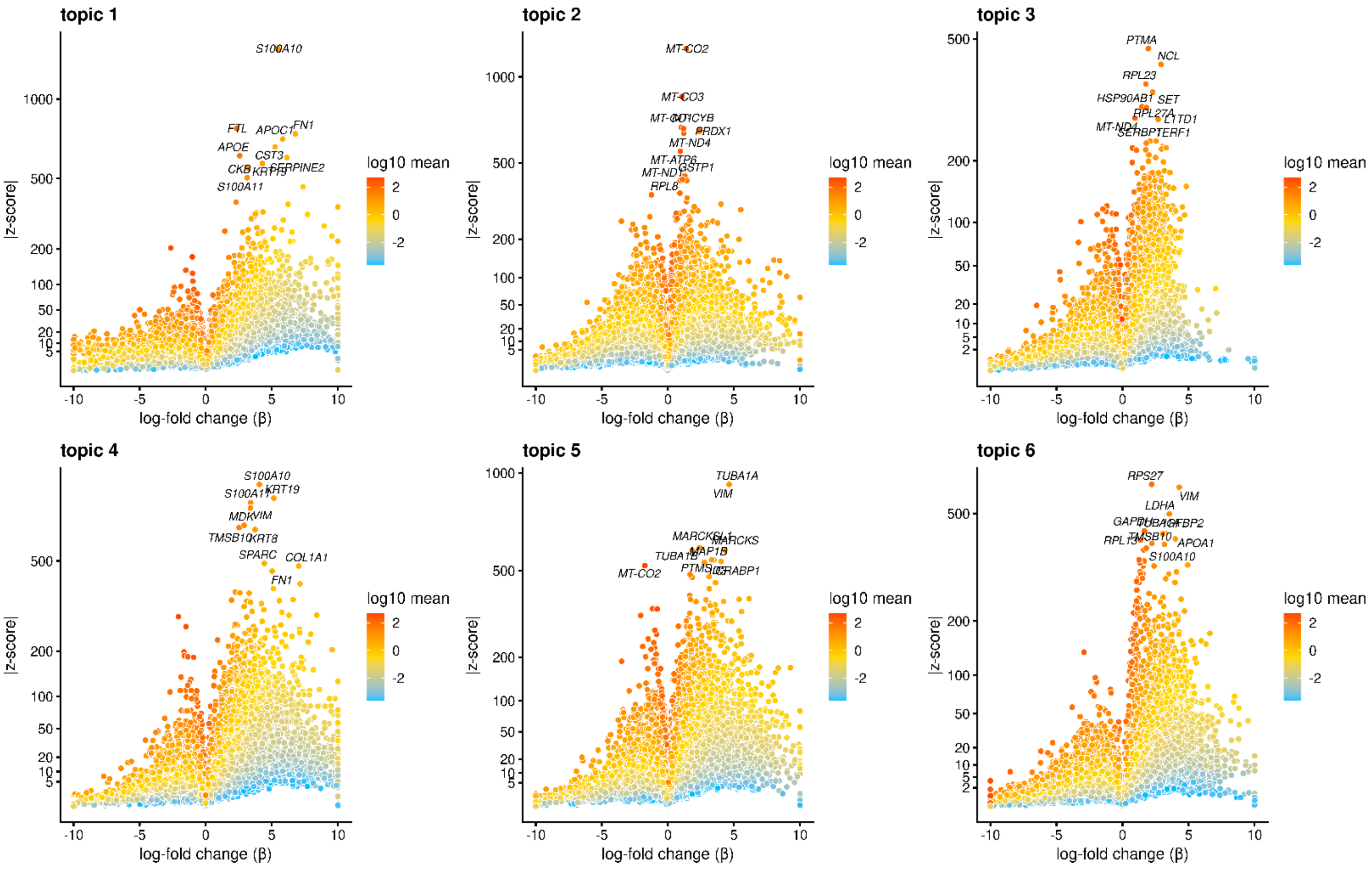


**Figure S6: Volcano Plot showing genes differentially expressed in each topic from the k=6 topic analysis.** Points are colored by the average count on the logarithmic scale. The top 10 driver genes of each topic are labelled.

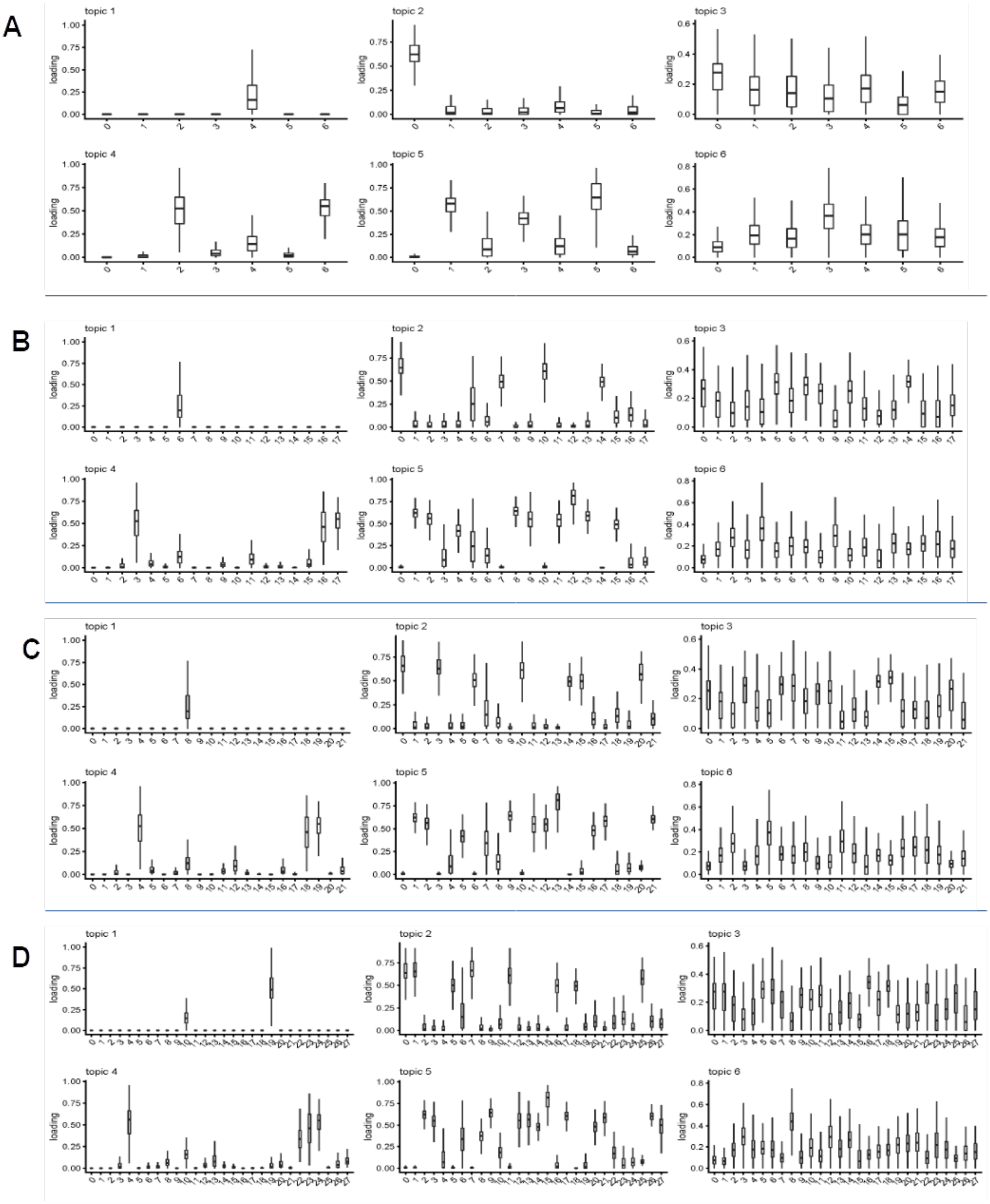


**Figure S7: Topic loadings on Seurat Clusters across clustering resolutions.** Barplots show the loading of each topic (from the k=6 analysis) on each seurat cluster at resolution 0.1 (A), resolution 0.5 (B), resolution 0.8 (C), and resolution 1 (D).

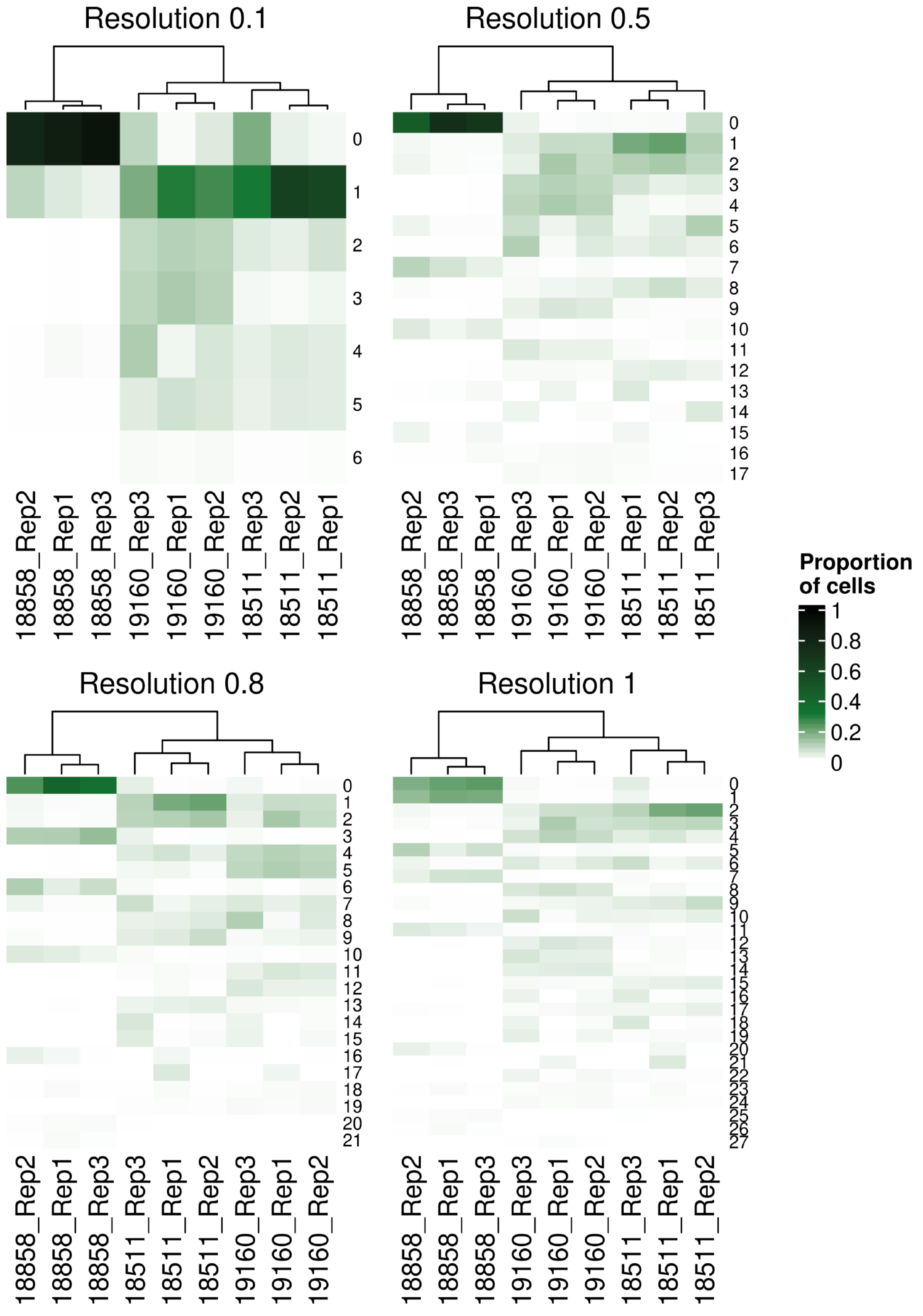


**Figure S8: Hierarchical clustering of samples individual-replicate groups by the proportions of cells from each group assigned to each Seurat cluster across clustering resolutions.**

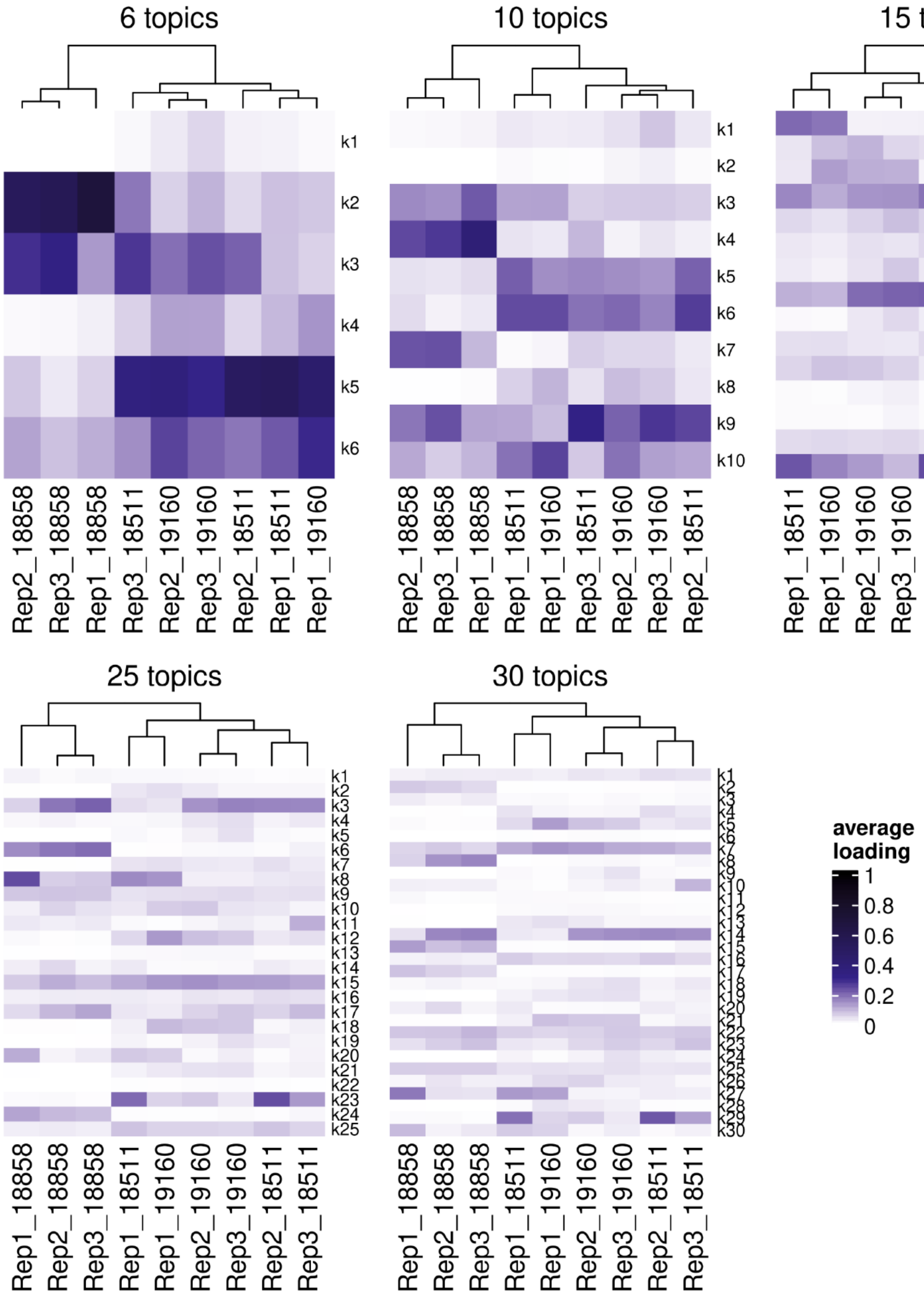


**Figure S9: Hierarchical clustering of samples individual-replicate groups by the loading of each topic with k=6, k=10, k=15, k=25, and k=30 topics.**

63

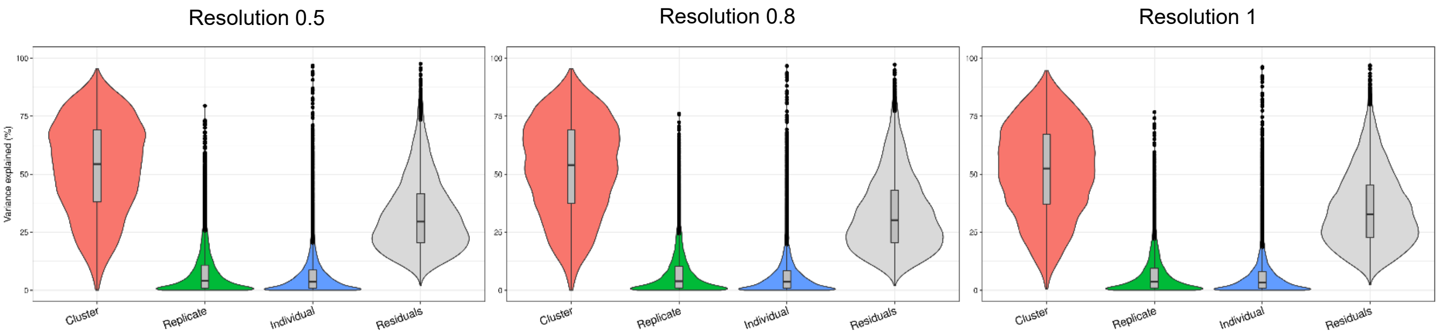


**Figure S10: Variance explained by biological and technical factors at higher clustering resolutions.** Violin plots showing the percent of variance in gene expression explained by cluster, replicate, and individual in this data set after partitioning variance in pseudobulk samples.

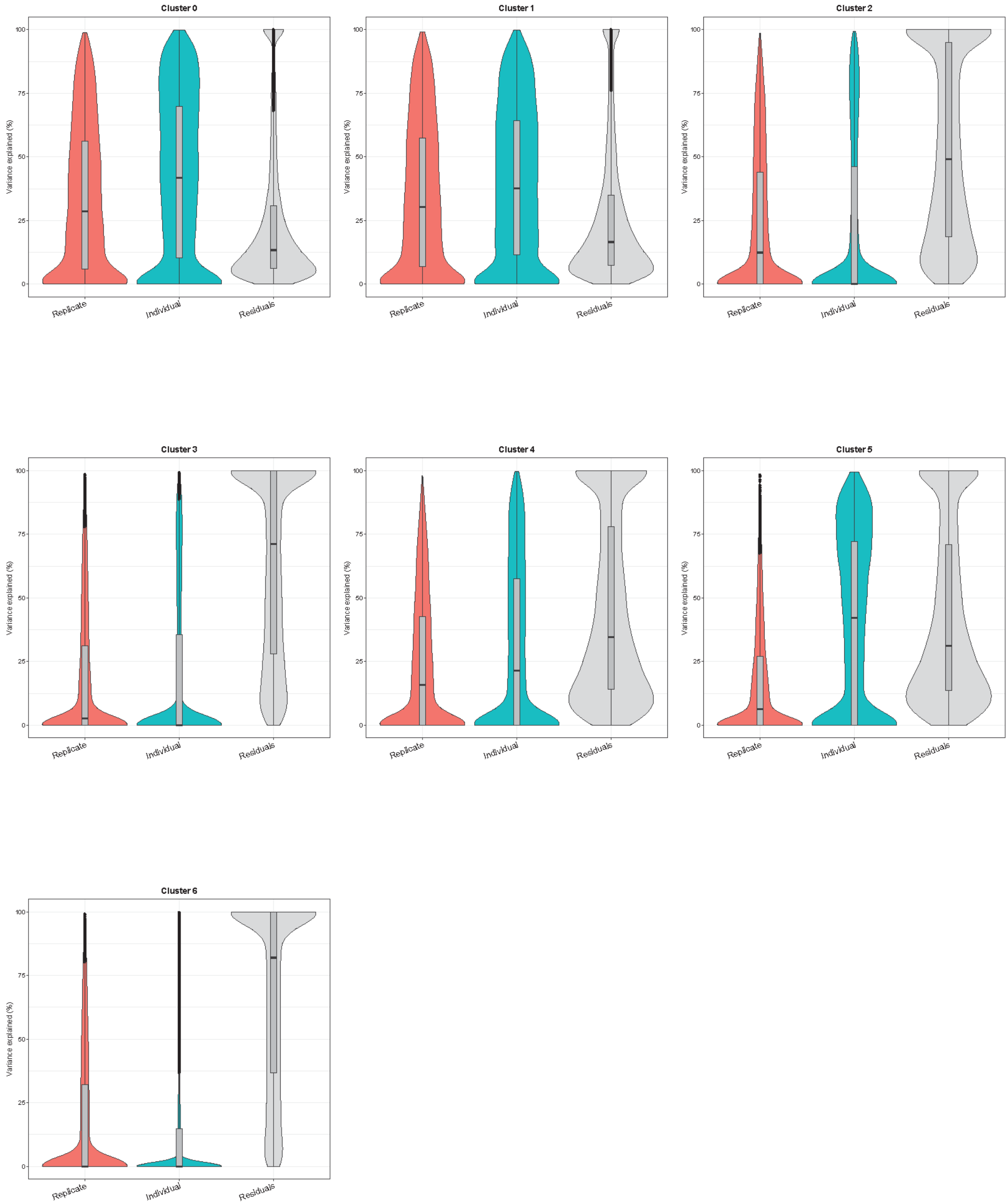


**Figure S11: Variance Partitioning by Seurat cluster using pseudobulk samples.** Violin plots showing the percent of variance in gene expression explained by replicate and individual in each Seurat cluster (clustering resolution 0.1).

66

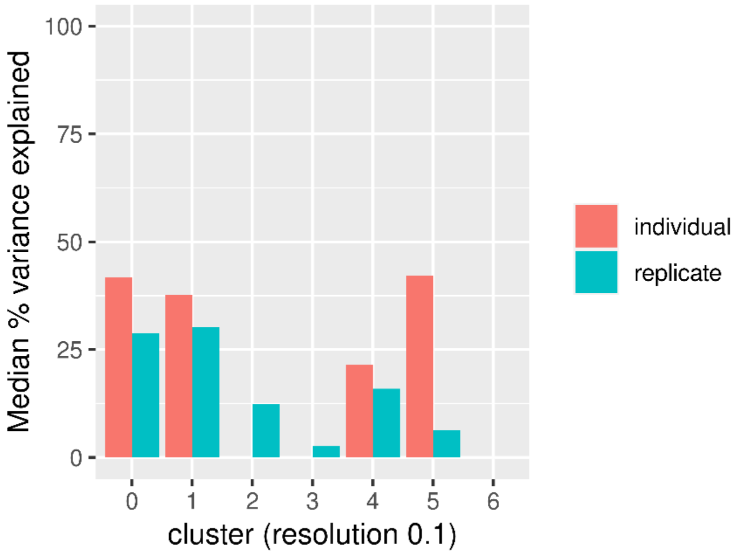


**Figure S12: Median PVE by replicate and individual in each cluster using pseudobulk samples.**

| | 0 | 1 | 2 | 3 | 4 | 5 | 6 |
|---|---|---|---|---|---|---|---|
| 18511_Rep1 | 99 | 2918 | 423 | 139 | 277 | 278 | 37 |
| 18511_Rep2 | 136 | 1958 | 35148 | 35 | 212 | 193 | 17 |
| 18511_Rep3 | 1508 | 2514 | 372 | 128 | 272 | 250 | 35 |
| 18858_Rep1 | 5995 | 526 | 29 | 28 | 116 | 35 | 3 |
| 18858_Rep2 | 4005 | 703 | 4 | 3 | 24 | 9 | 0 |
| 18858_Rep3 | 4882 | 235 | 5 | 9 | 47 | 12 | 0 |
| 19160_Rep1 | 81 | 2870 | 997 | 1129 | 196 | 634 | 63 |
| 19160_Rep2 | 179 | 1025 | 361 | 383 | 221 | 207 | 42 |
| 19160_Rep3 | 808 | 1634 | 747 | 819 | 1003 | 372 | 98 |

**Table S2: Number of cells per cluster (resolution 0.1) from each individual-replicate sample.**

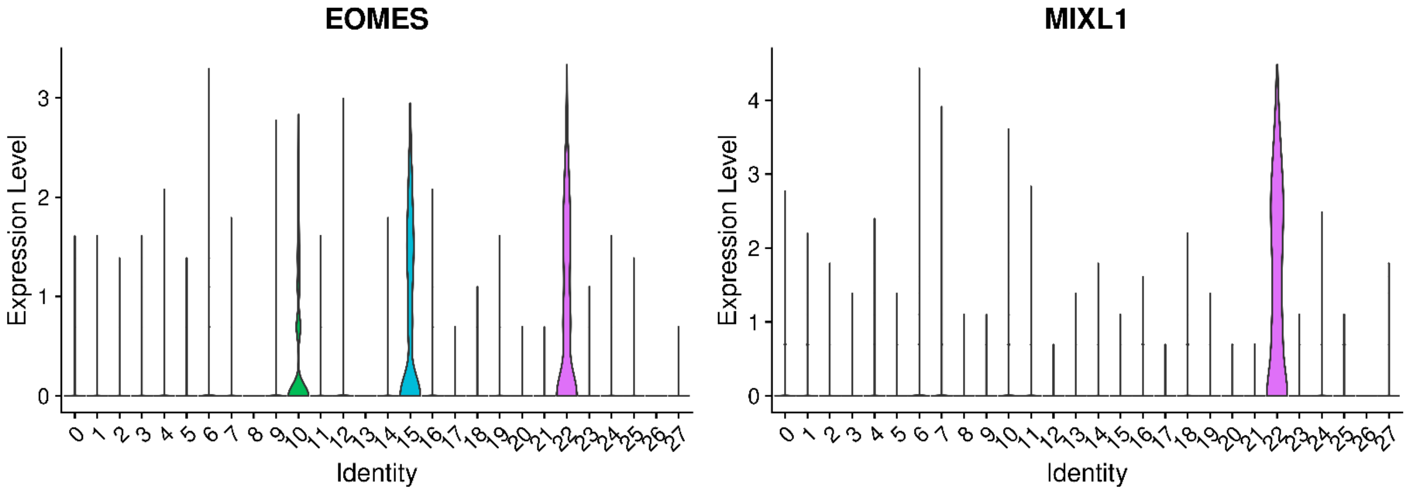


**Figure S13: Normalized expression of primitive streak markers (EOMES, MIXL1) in Seurat clusters defined at clustering resolution 1.**

68

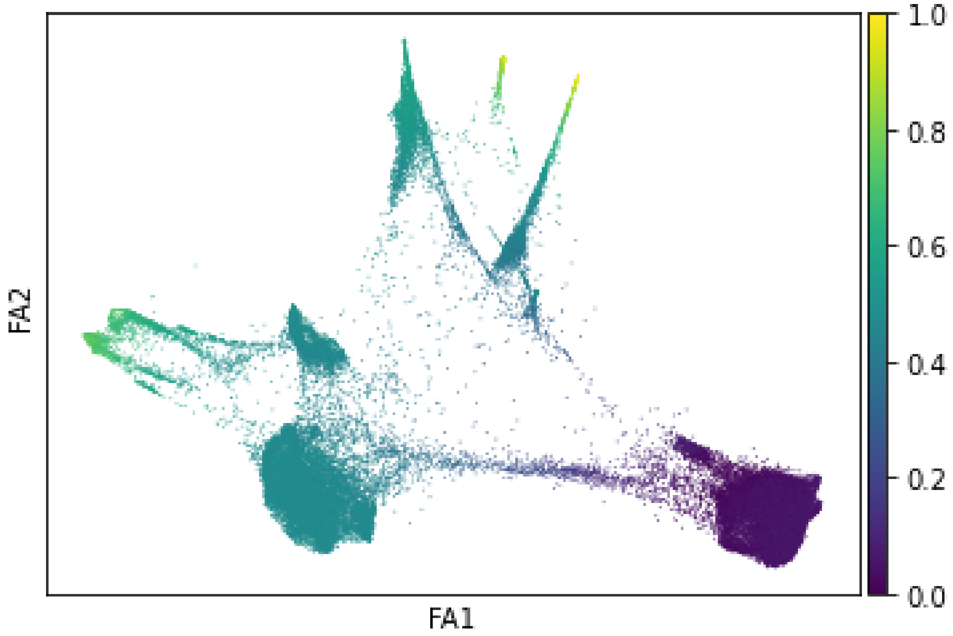


**Figure S14: Diffusion pseudotime values across EB cells visualized with force atlas.**

A

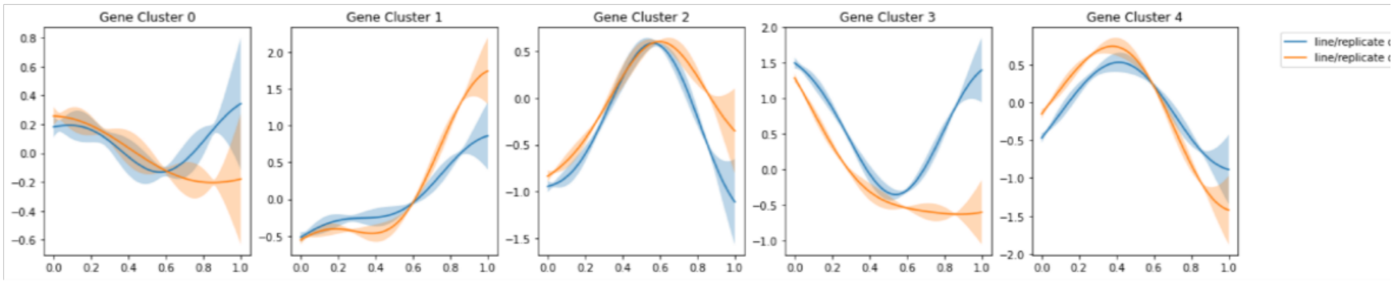


B

| | 3 bonferonni-adjusted | 4 bonferonni-adjusted |
|---|---|---|
| **HALLMARK_MITOTIC_SPINDLE** | 1 | 2.17277e-06 |
| **HALLMARK_G2M_CHECKPOINT** | 1 | 1.48348e-22 |
| **HALLMARK_ESTROGEN_RESPONSE_EARLY** | 0.000267846 | 1 |
| **HALLMARK_MTORC1_SIGNALING** | 6.99e-08 | 1 |
| **HALLMARK_E2F_TARGETS** | 1 | 3.49535e-25 |
| **HALLMARK_MYC_TARGETS_V1** | 0.192287 | 9.34926e-06 |
| **HALLMARK_MYC_TARGETS_V2** | 5.43658e-07 | 1 |
| **HALLMARK_OXIDATIVE_PHOSPHORYLATION** | 0.0026008 | 1 |
| **HALLMARK_REACTIVE_OXIGEN_SPECIES_PATHWAY** | 0.000274695 | 1 |

C

| | cell line/batch | assignment |
|---|---|---|
| **0** | SNG-NA18511--Batch1 | 2 |
| **1** | SNG-NA18511--Batch2 | 2 |
| **2** | SNG-NA18511--Batch3 | 2 |
| **3** | SNG-NA18858--Batch1 | 1 |
| **4** | SNG-NA18858--Batch2 | 2 |
| **5** | SNG-NA18858--Batch3 | 1 |
| **6** | SNG-NA19160--Batch1 | 2 |
| **7** | SNG-NA19160--Batch2 | 2 |
| **8** | SNG-NA19160--Batch3 | 2 |

**Figure S15: Cluster assignment by Split-GPM and gene set enrichment in the neuronal lineage.** A) Dynamic expression patterns of identified gene modules in each cluster of replicate-individual samples. B) Table showing bonferonni-adjusted p-values from gene set enrichment analysis of gene modules. C) splitGPM cluster assignments of each individual-batch sample based on shared patterns of dynamic gene expression.

A

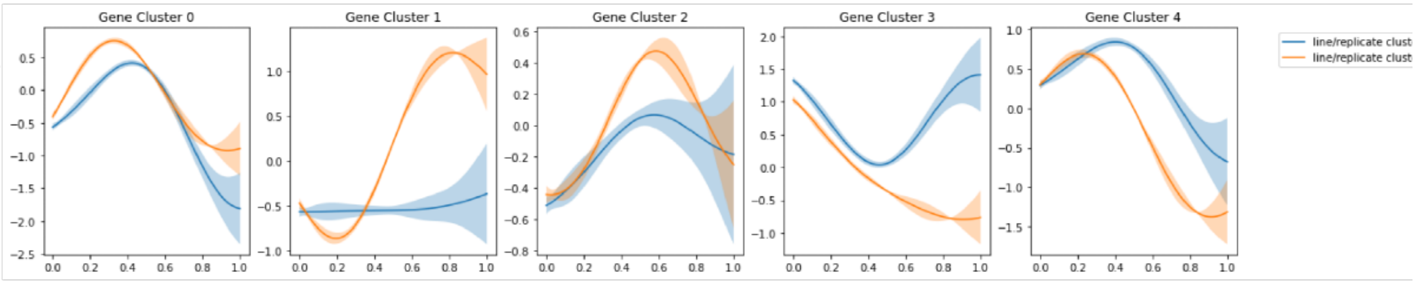


B

| | 1 | 4 |
|---|---|---|
| | bonferonni-adjusted | bonferonni-adjusted |
| **HALLMARK_CHOLESTEROL_HOMEOSTASIS** | 6.92618e-09 | 1 |
| **HALLMARK_MITOTIC_SPINDLE** | 1 | 4.92256e-13 |
| **HALLMARK_G2M_CHECKPOINT** | 1 | 1.17837e-27 |
| **HALLMARK_ADIPOGENESIS** | 4.37992e-05 | 1 |
| **HALLMARK_COMPLEMENT** | 7.02289e-06 | 1 |
| **HALLMARK_E2F_TARGETS** | 1 | 2.31674e-33 |
| **HALLMARK_MYC_TARGETS_V1** | 1 | 6.21581e-10 |
| **HALLMARK_XENOBIOTIC_METABOLISM** | 4.90832e-10 | 1 |
| **HALLMARK_FATTY_ACID_METABOLISM** | 4.23928e-09 | 1 |
| **HALLMARK_COAGULATION** | 8.51378e-14 | 1 |
| **HALLMARK_BILE_ACID_METABOLISM** | 2.19004e-06 | 1 |
| **HALLMARK_PEROXISOME** | 0.000565521 | 1 |

C

| | cell line/batch | assignment |
|---|---|---|
| **0** | SNG-NA18511--Batch1 | 2 |
| **1** | SNG-NA18511--Batch2 | 2 |
| **2** | SNG-NA18511--Batch3 | 2 |
| **3** | SNG-NA18858--Batch1 | 2 |
| **4** | SNG-NA18858--Batch2 | 1 |
| **5** | SNG-NA18858--Batch3 | 1 |
| **6** | SNG-NA19160--Batch1 | 2 |
| **7** | SNG-NA19160--Batch2 | 2 |
| **8** | SNG-NA19160--Batch3 | 2 |

**Figure S16: Cluster assignment by Split-GPM and gene set enrichment in the hepatic lineage.** A) Dynamic expression patterns of identified gene modules in each cluster of replicate-individual samples. B) Table showing bonferonni-adjusted p-values from gene set enrichment analysis of gene modules. C) splitGPM cluster assignments of each individual-batch sample based on shared patterns of dynamic gene expression.

A

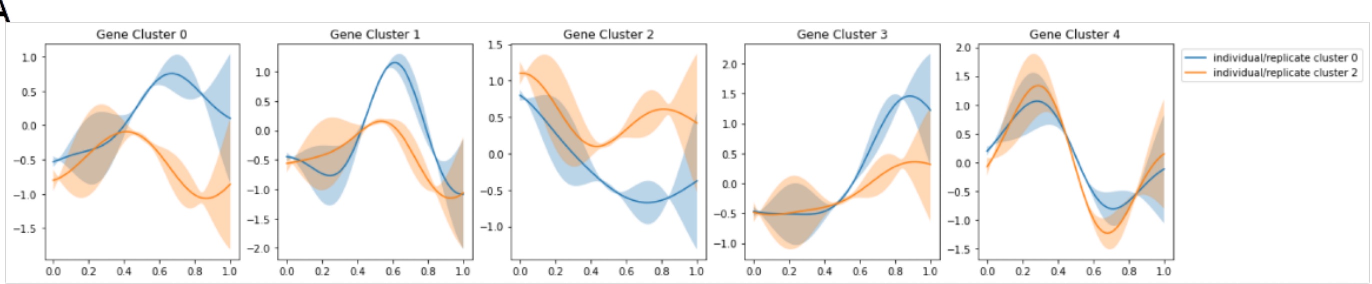


B

| | 1 | 2 | 3 | 4 |
|---|---|---|---|---|
| | bonferonni-adjusted | bonferonni-adjusted | bonferonni-adjusted | bonferonni-adjusted |
| HALLMARK_TNFA_SIGNALING_VIA_NFKB | 1 | 1 | 4.10533e-07 | 1 |
| HALLMARK_G2M_CHECKPOINT | 1 | 1 | 1 | 8.08289e-08 |
| HALLMARK_APOPTOSIS | 1 | 1 | 0.000118488 | 1 |
| HALLMARK_APICAL_JUNCTION | 1 | 1 | 0.000755343 | 1 |
| HALLMARK_COMPLEMENT | 1 | 1 | 6.5339e-06 | 1 |
| HALLMARK_MTORC1_SIGNALING | 1 | 1.78151e-05 | 1 | 1 |
| HALLMARK_E2F_TARGETS | 1 | 1 | 1 | 3.35097e-13 |
| HALLMARK_MYC_TARGETS_V2 | 1 | 0.000987191 | 1 | 1 |
| HALLMARK_EPITHELIAL_MESENCHYMAL_TRANSITION | 6.02819e-05 | 1 | 1 | 1 |
| HALLMARK_INFLAMMATORY_RESPONSE | 1 | 1 | 0.00462036 | 1 |
| HALLMARK_OXIDATIVE_PHOSPHORYLATION | 1 | 0.00108741 | 1 | 1 |
| HALLMARK_COAGULATION | 1 | 1 | 0.000313316 | 1 |
| HALLMARK_IL2_STAT5_SIGNALING | 1 | 1 | 0.00202719 | 1 |

C

| | cell line/batch | assignment |
|---|---|---|
| 0 | SNG-NA18511--Batch1 | 0 |
| 1 | SNG-NA18511--Batch2 | 0 |
| 2 | SNG-NA18511--Batch3 | 0 |
| 3 | SNG-NA18858--Batch1 | 2 |
| 4 | SNG-NA18858--Batch2 | 2 |
| 5 | SNG-NA18858--Batch3 | 2 |
| 6 | SNG-NA19160--Batch1 | 0 |
| 7 | SNG-NA19160--Batch2 | 0 |
| 8 | SNG-NA19160--Batch3 | 0 |

**Figure S17: Cluster assignment by Split-GPM and gene set enrichment in the endothelial lineage.** A) Dynamic expression patterns of identified gene modules in each cluster of replicate-individual samples. B) Table showing bonferonni-adjusted p-values from gene set enrichment analysis of gene modules. C) splitGPM cluster assignments of each individual-batch sample based on shared patterns of dynamic gene expression.

75

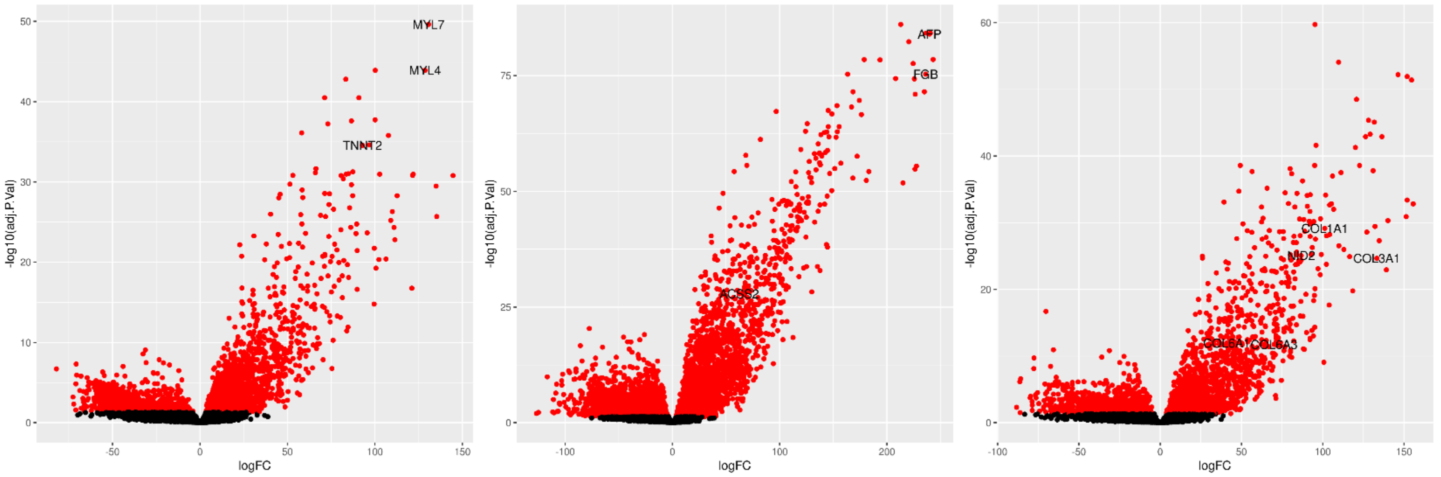


**Figure S18: Differential expression of known marker genes in reference annotated EB cell types.** Left) Volcano plot of DE genes in annotated cardiomyocytes compared to all other cell types with known cardiomyocyte marker genes labeled (MYL7, MYL4, TNNT2). Middle) Volcano plot of DE genes in annotated hepatoblasts compared to all other cell types with known hepatoblast marker genes labeled (AFP, FGB, ACSS2). Right) Volcano plot of DE genes in annotated mesothelial cells compared to all other cell types with known mesothelial marker genes labeled (NID2, COL1A1, COL6A3, COL3A1, COL6A1).

**Table S3:** Available as a .csv file. Contains the Limma differential expression results (logFC, AveExpr, t, P.Value, adj.P.Val, B) for all tested genes in each Seurat cluster when clusters were defined at resolution 0.1.

**Table S4:** Available as a .csv file. Contains the Limma differential expression results (logFC, AveExpr, t, P.Value, adj.P.Val, B) for all tested genes in each Seurat cluster when clusters were defined at resolution 0.5.

**Table S5:** Available as a .csv file. Contains the Limma differential expression results (logFC, AveExpr, t, P.Value, adj.P.Val, B) for all tested genes in each Seurat cluster when clusters were defined at resolution 0.8.

**Table S6:** Available as a .csv file. Contains the Limma differential expression results (logFC, AveExpr, t, P.Value, adj.P.Val, B) for all tested genes in each Seurat cluster when clusters were defined at resolution 1.
